# *Ratchetaxis* in channels: cells move directionally by pushing walls asymmetrically

**DOI:** 10.1101/2020.04.03.023051

**Authors:** Emilie Le Maout, Simon Lo Vecchio, Praveen Kumar Korla, Jim Jinn-Chyuan Sheu, Daniel Riveline

## Abstract

Cell motility is essential in a variety of biological phenomena ranging from early development to organ homeostasis and diseases. This phenomenon was so far mainly studied and characterized on flat surfaces in vitro whereas this situation is rarely seen in vivo. Recently, cell motion in 3D microfabricated channels was reported to be possible, and it was shown that confined cells push on walls. However, rules setting cell directions in this context were not characterized yet. Here, we show by using assays that ratchetaxis operates in 3D ratchets on fibroblasts and on epithelial cancerous cells. Open ratchets rectify cell motion, whereas closed ratchets impose a direct cell migration along channels set by the cell orientation at the channel entry point. We also show that nuclei are pressed at constrictions zones through mechanisms involving dynamic asymmetries of focal contacts, stress fibers, and intermediate filaments. Interestingly, cells do not pass these constricting zones when defective in the keratin fusion implicated in squamous cancer. By combining ratchetaxis with chemical gradients, we finally report that cells are sensitive to local asymmetries in confinement and that topological and chemical cues may be encoded differently by cells. Altogether our ratchet channels could mimic small blood vessels where cells are confined: cells would probe local asymmetries which would determine their entry into tissues and direction. Our results could shed light on invasions mechanisms in cancer.

## Introduction

Directed cell migration plays a key role in many physiological, pathological or developmental processes (1-4). Its dynamics has been widely and mainly studied far from physiological conditions, on 2D flat surfaces (5,6). However, in tissues or in capillaries, cells are often confined, and their cytoskeleton and organelles need to deform and to reshape in order to pass obstacles and sustain motion.

To test these mechanisms, it has been shown in microchannels that motility in confined cells could be distinct from the motility acquired on a flat surface: cells “push” on walls (7-9). And it was reported that large organelles such as nuclei undergo important deformations with biological consequences (10-11). Nevertheless, it is not clear yet how the cell cytoskeleton reorganizes to face local asymmetries in the channels or in squeezing zones. These types of situations can be found in small veins and in venules where microscopic venous valves (MVV) are present (12) introducing ‘bottlenecks’ in the migratory path. In addition, in some squamous cancers, cells are known to be defective for keratin, an intermediate filament essential to shape the scaffold surrounding the nucleus (13). Defects in these structures and their implications in motility in confined spaces have not been reported. It is also not clear if the broken symmetry given by bottlenecks could direct cell motion as it has been shown previously with other configurations by *ratchetaxis* (14-20). In this work, we show that periodic asymmetries direct cell motion when cells are confined in open channels. When totally confined, direction of migration is exclusively set by the cell entry point in channel: cells cannot repolarize and ‘ignore’ cues provided by the geometry.

To characterize the cellular mechanisms involved in this directed motion, we next studied the dynamics of actin, focal contacts, the nucleus and also the keratin array surrounding the nucleus when cells pass through bottlenecks. We report anisotropies in distributions of cytoskeletal proteins with actin bundles located at the bottom and the top of the cell between which the nucleus can be deformed. Focal contacts are mainly located at the channel wall specifically at the junctions between ratchet units. The keratin array also exhibits an asymmetric distribution where keratin accumulates at the rear of the nucleus, potentially to facilitate deformation. Finally, we combined chemical gradient with the channel experiment and we report that cells are able to sustain their migration when the gradient is removed only when the direction of anisotropies is not an obstacle; otherwise, cells stop migrating. Altogether, our results describe how cells can migrate and deform through confined spaces, providing potentially new ways to envision cell motility in small blood capillaries *in vivo*.

## Results

In order to mimic blockage of cells in small veins, we designed microfabricated channels with periodic asymmetries to bias polarity and rectify cell motions through *ratchetaxis*. To design channels, we tested different conditions with various bottlenecks sizes and angles. For strong confinements, cells were not able to move, and otherwise nuclei were less ‘squeezed’ (Figure S1). The best balance between optimal confinement and sustained migration was obtained for the condition ‘a16i4’ where a corresponds to the tip angle and i to the opening width.

Two sets of configurations were considered (Figure 1a) to evaluate potential effects of full confinement: open channels, when cells are only confined by sides (Movies S1 and S2), and closed channels where cells are totally confined (Movies S3 and S4). For each condition, ratchets were compared with linear channels which dimension was taken by the width I of the ratchet constriction (see Figure 1b bottom).

**Figure 1.**
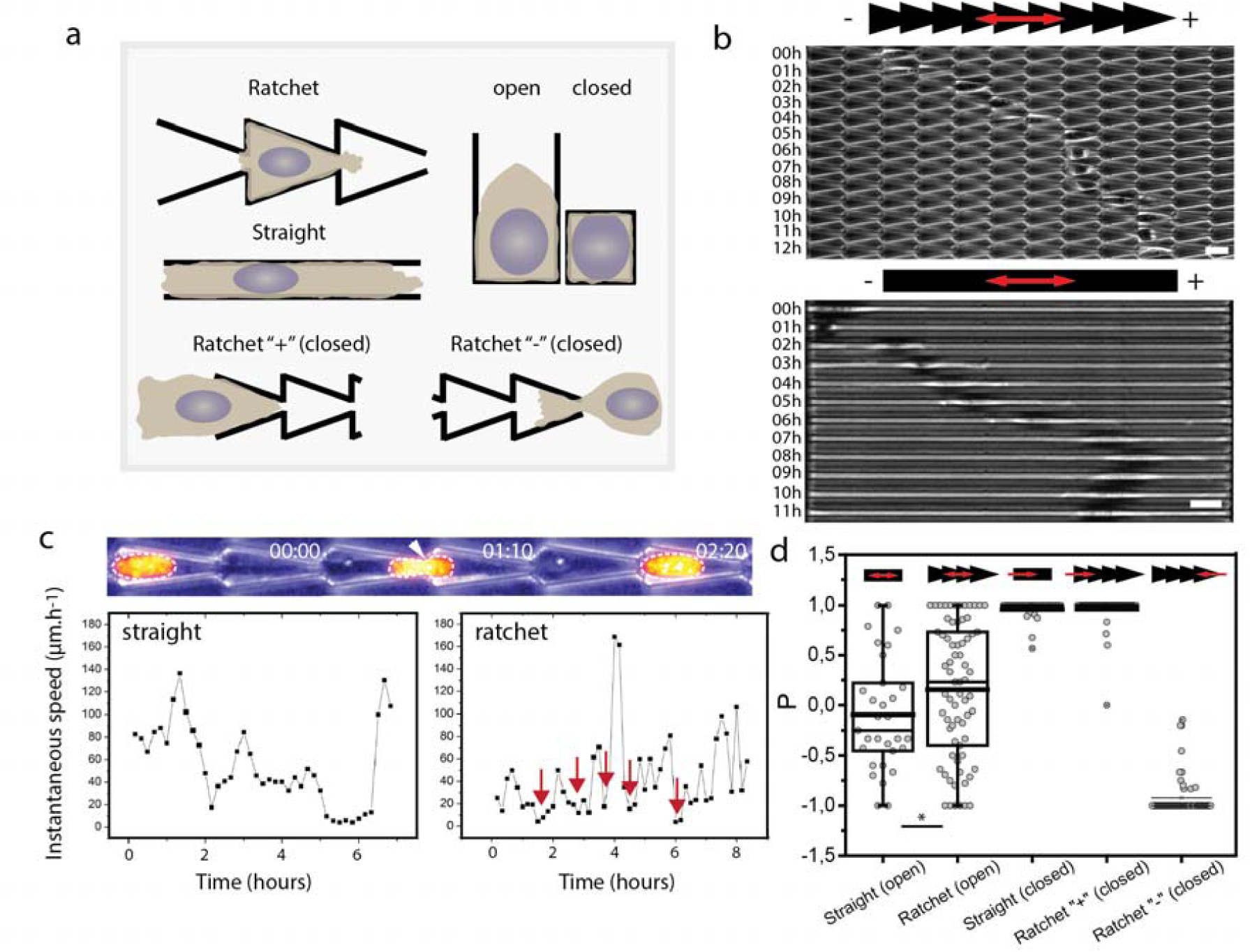
Cell motility in 3D ratchets. (a) Configurations tested in this study. Two types of confinements are produced: side-only confinement with open channels and full confinement with closed channels. Ratchet are defined as “+” when cells are directed towards the tip of the ratchet and “-” when cells are directed against the tip of the ratchet. (b) Cells migrate in ratchets and in straight channels. Here open channels are shown. Scale bars: 20µm. (c) Nucleus has to deform to pass the bottleneck in closed configurations, leading to drops in velocities corresponding to pauses (red arrows). In straight channels, velocity is almost always non-zero and migration is facilitated. (d) Mean bias per step *p* as a function of the tested condition. In open configurations, cells migrate preferentially in the “+” direction, consistent with previous 2D works (12-18). In closed channels, cells are totally biased by the entry point with no change in direction resulting in distributions peaked at 1 and −1. Significance has been tested with distributions comparison using a Kolmorogov test.

NIH3T3 cells were able to move along the channels over the whole experimental time (12h) (Figure 1b and Movies S1-S4). As in *ratchetaxis* on topographical motifs, the nucleus seems to be the large organelle preventing passage, as assessed by pausing phases in motion recorded in the ratchet, and absent when cells move along linear channels (Figure 1c). Deformation of the nucleus was clearly visible in our experiments (Figure 1c and Movie S5) and for different cell types (Figure 2e and Figure S2).

**Figure 2.**
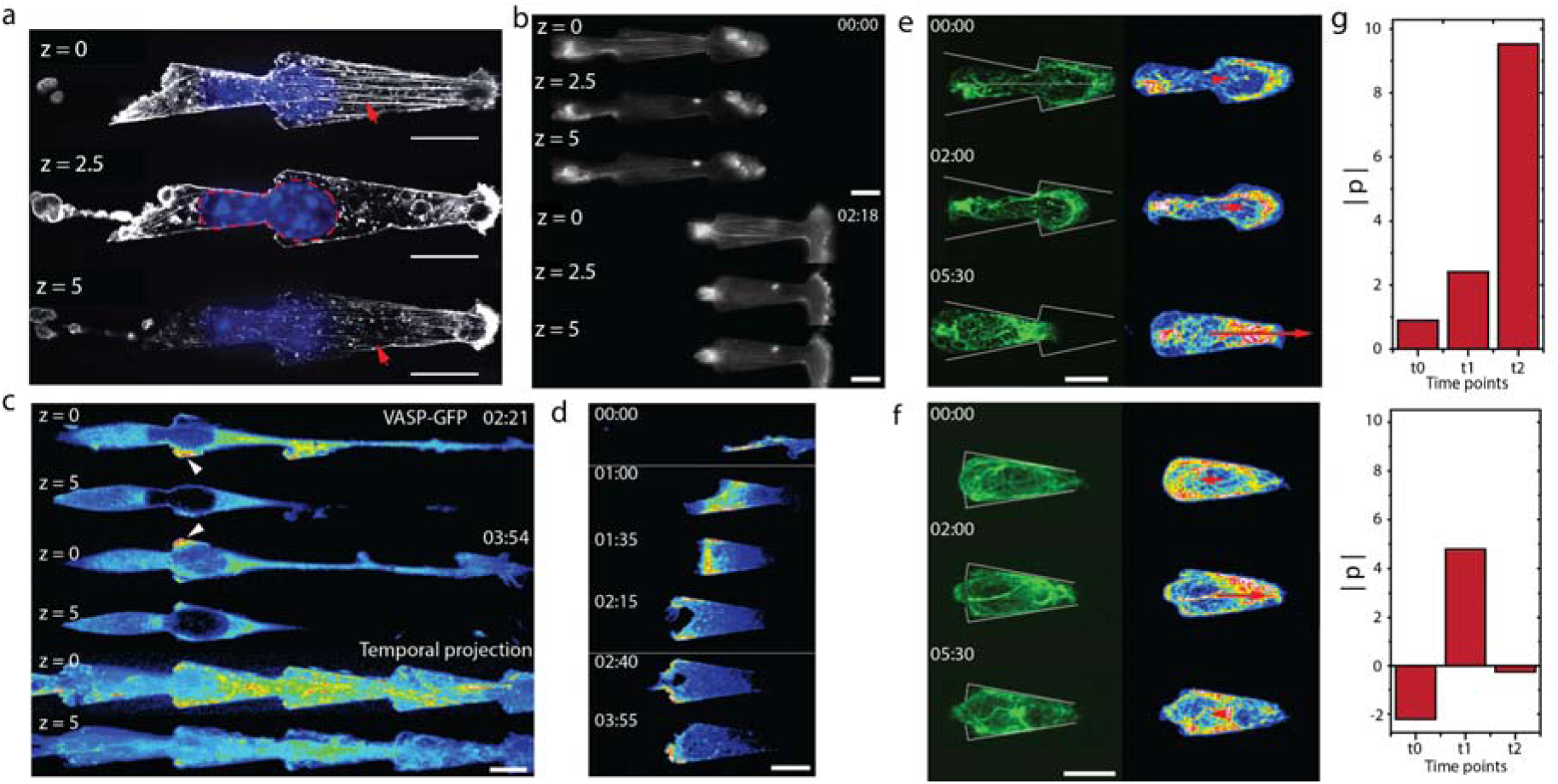
Anisotropies in cytoskeleton correlate with nucleus deformation. (a) NIH3T3 in closed ratchet. Staining shows actin and nucleus. Arrows indicate actin bundles at the top and at the bottom, surrounding the nucleus deformed at the bottleneck. (b) Lifeact-mcherry live. Actin bundles are present during the whole motion of cell. (c) VASP-GFP in NIH3T3. Focal contacts are mainly present at the bottleneck-bottom and also in protrusions sent by cell. (d) Protrusion growth with VASP-GFP. Focal contacts mainly nucleate at the interface between the wall and the surface. (e) K14-GFP showing the keratin cytoskeleton in CAL27 oral squamous cells with its anisotropy map. Keratin is distributed along the rear of the nucleus during its passage through the bottleneck. (f) CAL27 expressing modified keratin fusion protein with its anisotropy map. (g) Quantification of the norm of polarization 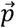 for WT K14-GFP (up) and for Keratin fusion (bottom). Scale bar = 15µm in all the panels. Time in hh:mm.

In the open channel configuration, cells were rectified along the direction of the ratchet (Movie S1), consistent with the two other configurations probed before (14-17). This is shown with the mean displacement vector (Figure S3). The bias is also characterized with the measurement of the mean bias per step, p, defined as (18):

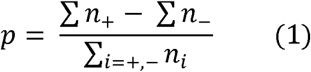

where n_i_ corresponds to a step (in lattice unit) made in the i direction, and where i = + if the cell moves towards the tip of the ratchet, and i = - if the cell moves towards the basis of the ratchet. This parameter provides a clear read-out of the bias in cell migration and puts forward a significant difference between cell motion in ratchets and straight channels in open configurations (Figure 1d).

Surprisingly, cells that are fully confined exhibit a persistent migration depending only on the *broken symmetry of the entry point* with no sensitivity to the ratchet orientation afterwards (Movies S3 and S4). It appears that up to the channel length probed in our devices, cells keep their motion throughout the channels with both directions leading to distributions of p peaked around 1 and −1 depending on the entry point position (Figure 1d). This result suggests that the entry might be critical when transposed to *in vivo* situations: when cells enter a capillary sufficiently small, repolarization would be impossible and direction of motion would be kept – and in the absence of chemical gradient.

Next, we focused on closed configurations viewing this set up as the closest to *in vivo* situations. We asked how cells could pass bottlenecks of ratchets. We imaged the localizations and dynamics of actin cytoskeleton, intermediate filaments and focal contacts. Stress fibers were present below and above the nucleus, like in the topographical ratchets (Movie S6, Figure 2a-b and (17)). Focal contacts were observed as well (Movie S7 and Figure 2c-d). However, they specifically accumulated right after the bottleneck of ratchet (Figure 2c-d). Distributions of these markers are established hallmarks for local stress generation. This suggests that cells anchor and ‘pull’ locally to deform the nucleus which allows the cell passage. Sequences of such correlations between cellular structures and cell motion are reported in Figure 2 and in Movies S6-S7.

Defects in keratin cytoskeleton have been reported to be involved in some squamous cancers (13) but no dynamics analysis and migratory assays were reported so far. To determine whether defects in keratin impair motion in confined spaces, we used oral squamous epithelial cells CAL27, expressing wild type keratin tagged with GFP. We also used a CAL27 mutant expressing a fusion keratin protein involved in squamous cancers. For both cell lines, no net motion was recorded on flat coverslips as expected for epithelial cells. However, WT cells could enter in closed ratchets and migrate efficiently therefore confirming the difference in migratory behaviors between flat and 3D environments. Strikingly, in these cells, keratin was a dynamic structure with strong anisotropies in distribution (see Movie S8 and Figure 2e). Accumulation of keratin was visible at the rear of the nucleus when the cell passed through the bottleneck (Figure 2e). This is quantified by the measurement of polarity as a readout of anisotropies in keratin distributions (see Figure 2e-g). However, in the mutant strains expressing keratin fusion, keratin network was isotropic (Figure 2f-g). Interestingly, these cells were arrested at the bottleneck and did not move to the next ratchet unit (Movie S9 and Figure S4a). We checked that nuclei had similar dimensions in wild type situations and in mutants (Figure S4b), which suggests that defects in intermediate filaments are correlated to the arrest. Interestingly, fusion cell line had no – or very small – structured keratin array surrounding the nucleus and the nucleus was frequently fragmented (Figure S4a). This suggests that dynamics of the keratin cytoskeleton is also important to deform and maintain integrity of large organelles such as nuclei and this sheds light to the potential involvement of intermediate filaments in the dynamic remodeling of cellular organelles. The fact that this fusion is reported in squamous cancer suggests that cells could be blocked within small capillaries allowing them to probe the endothelial layers and potentially penetrate tissues (21-22).

We next combined motion in closed channels with *chemotaxis* to see the potential interplay between topological and chemical cues. We designed assays where cells face a serum gradient 0% to 10% (see Methods). The gradient and the slope of the gradient was quantified with the visualization of TRITC-dextran incorporated in the solution (Figure 3a). In both ratchet orientations, cells migrate towards the high serum concentration (Figure 3b). However, upon removal of the chemical gradient, an unexpected effect appeared: cells facing the ratchet direction could move while cells in the opposite direction did not (Movies S10-11 and Figure 3c-d). This suggests that polarity induced by gradient could be distinct from the polarity set by topography. We quantified the observed phenomenon: 2 hours after gradient removal, cells in the ratchet direction (+) move on average twice more than the cells facing the opposing direction (-) (Figure 3e). When we align all the trajectories sequences with respect to the gradient removal, we observe that the ratio of velocities for each case 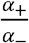 is about 10 confirming this unexpected result (Figure 3f).

**Figure 3.**
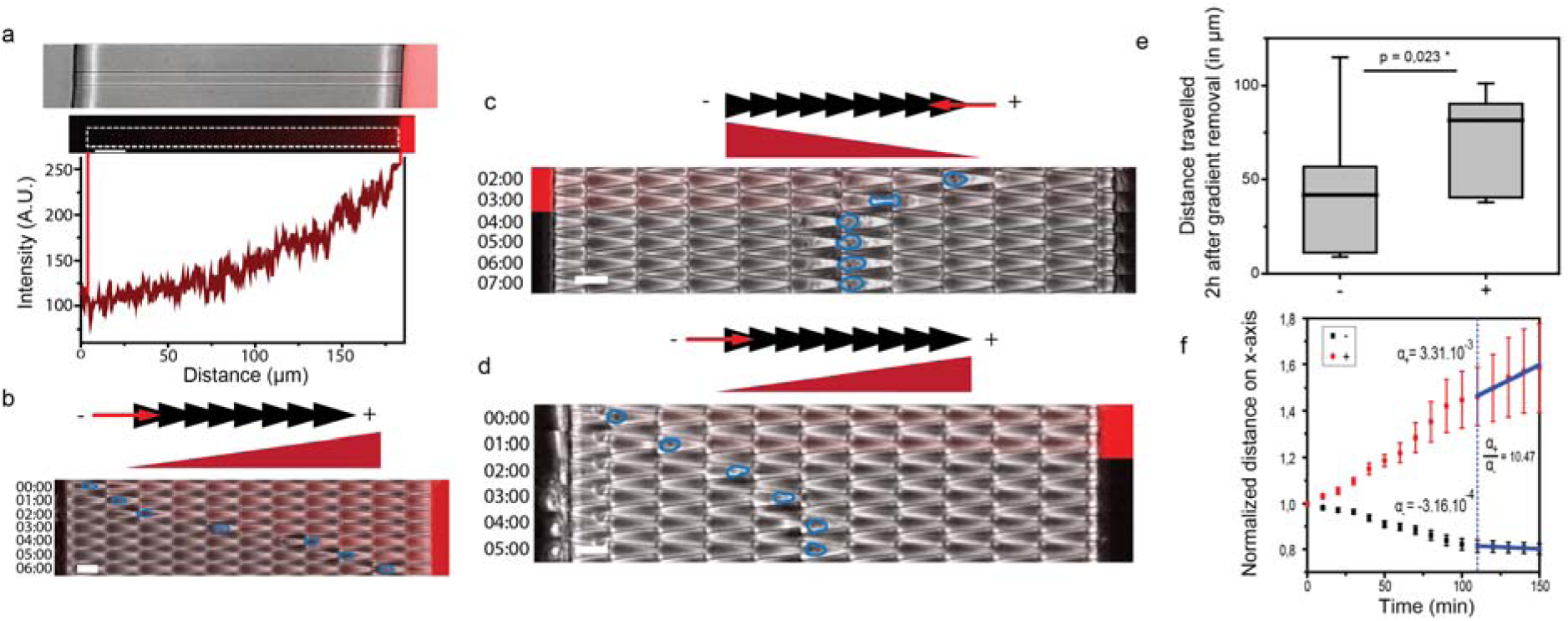
Combination of *ratchetaxis* and chemotaxis. (a) Phase-contrast image of a straight channel (closed) with a serum gradient. The gradient is visualized with TRITC-dextran and the slope can be extracted. (b) NIH3T3 moving towards the high serum concentration. Contour of the nucleus is shown in blue. (c) After gradient removal, the cell stops its motion when it moves against the ratchet direction. When the cell faces the ratchet orientation (d), it continues to move after gradient removal. (e) Quantification of the distance travelled 2h after gradient removal. (f) Cells positions on the x-axis aligned with removal of gradient (dashed blue line). Slopes are extracted from the linear fits after gradient removal and respective coefficients are displayed on the plot. Cells in the orientation of the ratchet (+) can migrate whereas cells in the opposite direction stop (-). Scale bar = 20µm in all the panels. Time in hh:mm.

## Discussion

Our results show that cells direction in ratchet channels has two types of behaviours. In open configurations, rectification occurs along the direction of the broken symmetry. In the closed configuration, direction is set by the polarization induced by the entry of cells in channels. Focal contacts, actin, and keratin anisotropy are involved in the motion, in particular to allow the passage of nuclei blocked at bottlenecks along other reports deciphering molecular interplays between the cytoskeleton and the nuclear envelop (23-24). Defects in keratin impairs motion. Finally, chemical gradients can induce directed motion, and cells become sensitive to the asymmetry of confinement when the chemical gradient is removed. Altogether, these rules open a variety of simple rules for understanding cell motion in confined situations.

Extrapolation of these results to *in vivo* cases is attractive. The facts that capillaries of the cell size are reported (12) and that cell blockage is established in vessels suggest that confinement could be an important phenomenon in physiological situations. From this respect, the entry of cells in channel and their subsequent motion could constitute a relevant assay to reproduce *in vitro* this situation. Single cells may probe their local environment and the entry in a vessel could be related to the polarization rules reported in our study. This could involve topographical asymmetry where cells would probe the local environment and/or exposure to changing chemical gradients which may bias cells to move directionally. Future works could test this hypothesis in ratchet configuration’s and in cellular systems prone to be responding to chemical gradients in cancer.

Our results with the fusion keratin are interesting along this line on its implication in cancer. Cells could not pass bottlenecks anymore, and this was associated to defects in the keratin network. Since this fusion was reported in medical cases, this suggests that an actual cancer situation is responding to confinement. And because initial arrest in the capillary is critical for tumor cells to metastasize to the second sites in distant organs, blockage of *ratchetaxis* by fusion keratin may provide advantages for tumor seeding, survival and proliferation (21-22). Impaired migration addressed here would be distinct from cell entry from primary tumors into blood vessels in which protease-mediated matrix remodeling play more roles.

The gradient experiment was puzzling. There seems to be an increase sensitivity related to the exposure to chemical gradients. Polarity cues in the cortex may lead to a local increase in surface tension. As a result, subsequent motion in the presence of an obstacle could be affected depending on the direction of the topographical anisotropy. These experiments suggest that distinct pathways may be triggered by chemical and by mechanical gradients.

Competition and cooperation were reported between different types of gradients and they could lead to compensation or increased motion. In this current study, we report that rules have to be revisited in closed configuration. This suggests that repolarization is possible in open configurations and in other reported cases only when cells can reorganize their polarity. There could be a locking mechanism of cell polarity which would be forcing cells in closed confinement to keep the same cues for long distance and time.

Finally, this study suggests that channels may be more relevant to study invasion than standard tests with Boyden chambers for example. In this latter case, chemical gradients are placed between two reservoirs, and the passage of cells – fibroblasts, epithelial cells for example – is tested through thin ‘cavities’ with a fraction of a cell size as dimension. Such configuration seems far from *in vivo* cases. Future studies could take this long and ratchet channel strategy to identify signaling networks modification in the context of cancer, along the strategy reported in this study for squamous cancer cell lines and their behaviours in closed ratchets.

## Acknowledgments

We thank J. Comelles, E. Pencreach, O. Pertz, P. Rossolillo, the Imaging Platform of IGBMC, the Riveline Lab. for help, constructs and discussions. We acknowledge funding from Région Grand Est, Alsace contre le cancer, the University of Strasbourg and the CNRS. This study with the reference ANR-10-LABX-0030-INRT has been also supported by a French state fund through the Agence Nationale de la Recherche under the frame programme Investissements d’Avenir labelled ANR-10-IDEX-0002-02.

## Figures and Movies Captions

**Figure S1.**
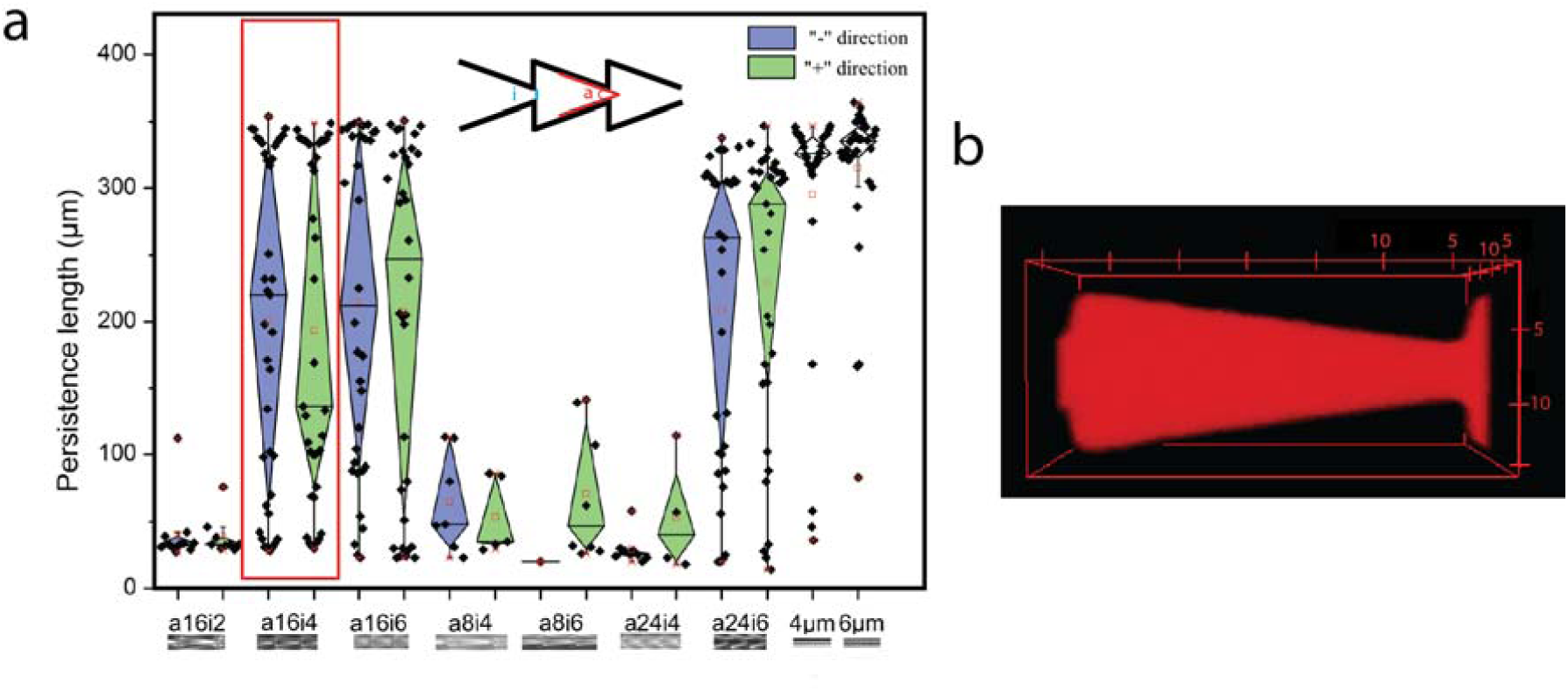
Comparison of different ratchet designs. (a) Distribution of persistence lengths for different closed ratchet configurations. The design used in the study is highlighted in red; straight channels are tested for comparison (two last conditions). (b) Visualization of the ratchet lattice unit corresponding to the configuration a16i4. The channel is filled with TRITC-dextran and imaged. Scale in µm.

**Figure S2.**
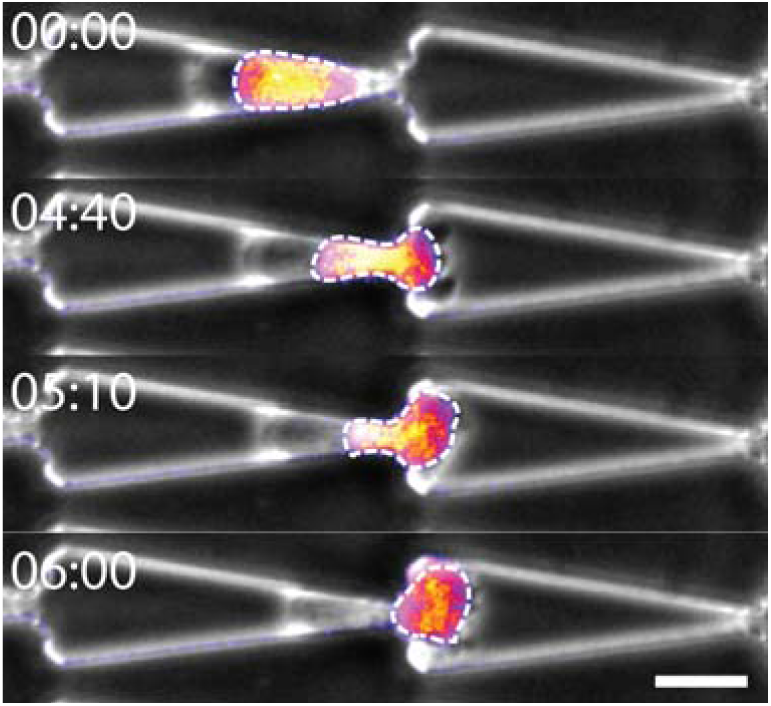
Deformation of nucleus for a HL60 cell moving in a ratchet (closed). Nucleus is outlined; label DAPI, Scale bar = 20µm. Time in hh:mm.

**Figure S3.**
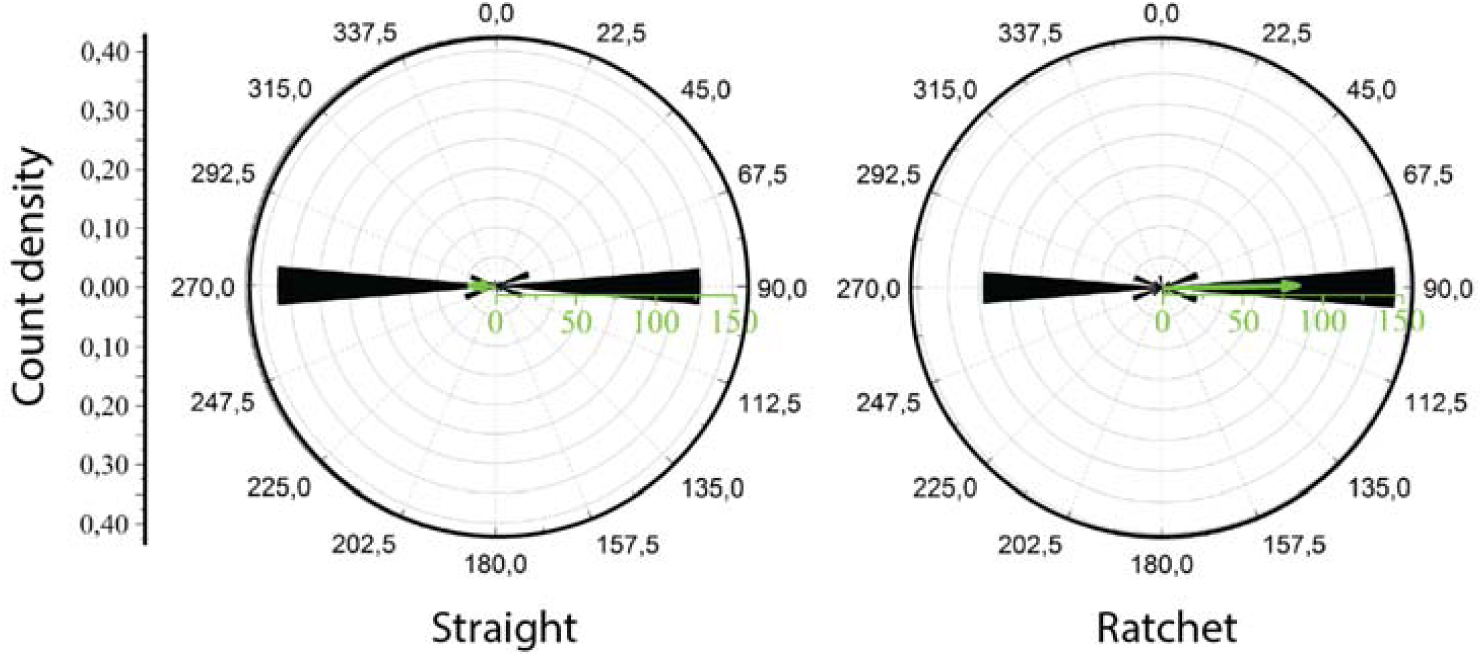
Mean displacement vector for straight channels and ratchets (open). All cell trajectories are averaged out to determine the mean displacement vector for each condition (see (17)).

**Figure S4.**
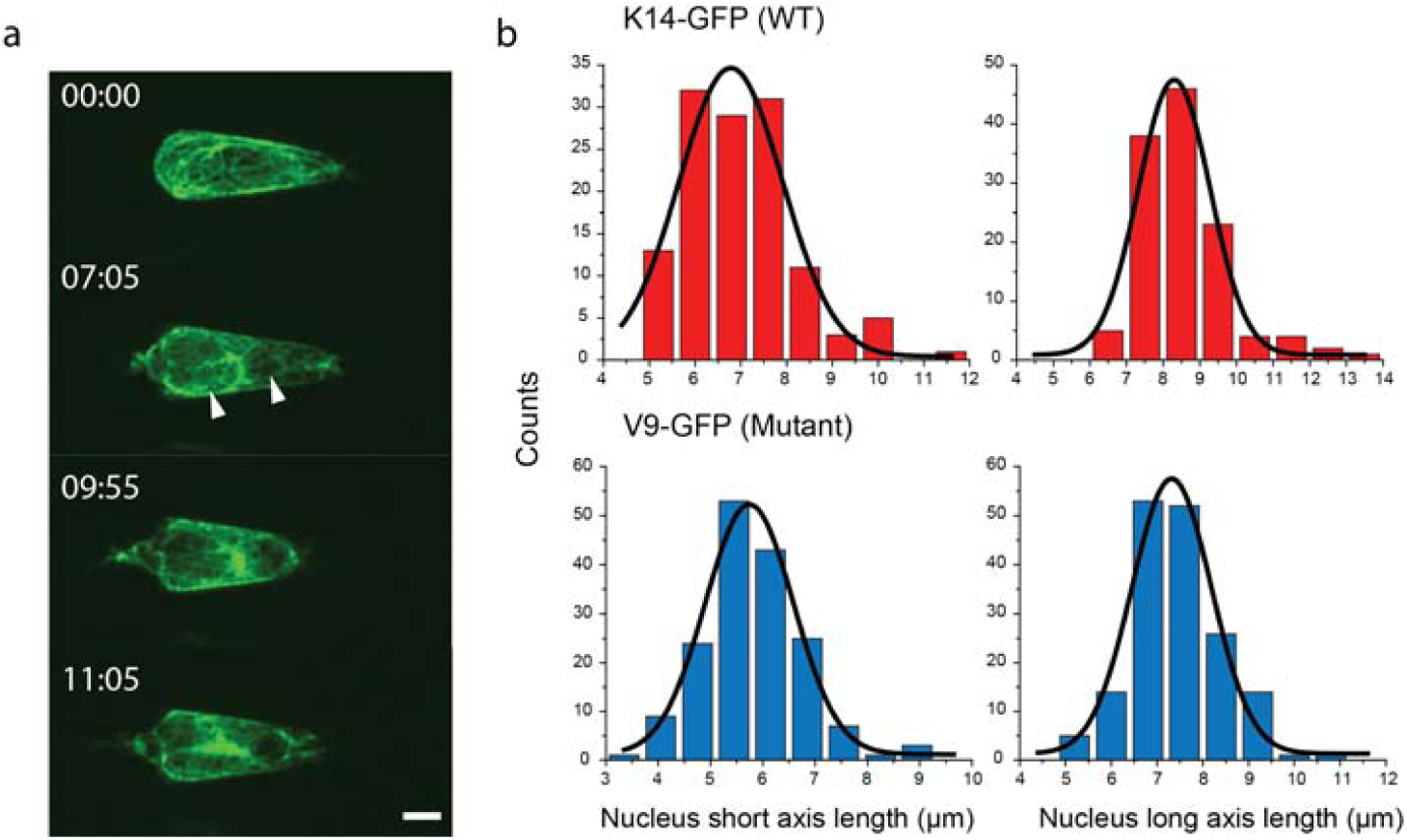
Fusion keratin (V7-GFP) mutant cells. (a) Cell inside the ratchet: motion to the next lattice unit is absent and the keratin array is ill-organized. Arrows indicate fragmentation of the nucleus. Time in hh:mm and scale bar = 5µm. (b) Distributions of nuclei sizes for V7 and K14-GFP. Nucleus dimensions are similar.

**Movie 1**. NIH3T3 cell in open ratchet, time in hh:mm, scale bar = 15µm

**Movie 2**. NIH3T3 cell in open straight channel, time in hh:mm, scale bar = 15µm

**Movie 3**. NIH3T3 cell moving along the direction ratchet (closed), time in hh:mm, scale bar = 15µm

**Movie 4**. NIH3T3 cell moving against the direction ratchet (closed), time in hh:mm, scale bar = 15µm

**Movie 5**. Nucleus deformation during passage, time in hh:mm, scale bar = 15µm

**Movie 6**. NIH3T3 actin distribution during motion. Z planes in µm from the bottom surface z=0. Time in hh:mm, scale bar = 10µm

**Movie 7**. NIH3T3 focal contact distribution during motion. Z = 0 (up) and upper plane z = 5µm (bottom). Time in hh:mm, scale bar = 10µm

**Movie 8**. Reorganization of keratin during passage, time in hh:mm, scale bar = 10µm

**Movie 9**. Fusion keratin cell blocked by the ratchet, time in hh:mm, scale bar = 10µm

**Movie 10**. NIH3T3 cell passing after removal of serum gradient (visible in red on the left when present), time in hh:mm, scale bar = 20µm

**Movie 11**. NIH3T3 cell blocked after removal of serum gradient (visible in red on the right when present), time in hh:mm, scale bar = 20µm

