## Supplementary material for "*Ratchetaxis* in channels: cells move directionally by pushing walls asymmetrically": Material and Methods

Materials and methods

Detailed protocols are available in the reference (25) with troubleshooting. Here, we briefly describe the main steps.

Microfabrication

SU-8 molds are prepared using standard UV-photolithography (25). Polydimethylsiloxane (PDMS, 1:10 w/w cross-linker:pre-polymer) (Sylgard 184 kit, Dow Corning, cat. DC184-1.1) is then poured and cured at 65°C for at least 4 hours.
Next, PDMS chips and glass coverslips are treated with oxygen plasma and bound together in two different ways depending on the configuration tested (closed or open) (25). Micro-channels chip is facing up for open channels while it is facing down for the closed configuration.

Cell culture and migration assays

NIH3T3 (ATCC) cells were culture in Dulbecco’s Modified Eagle’s Medium (DMEM) high glucose (Fisher Scientific, cat. 11574486) with 10% Bovine Calf Serum (BCS) (Sigma, cat. 12133C) and 1% Penicillin-Streptomycin.
HL60 cells were grown in suspension in RPMI 1640 (Gibco, cat. 22409031) supplemented with 40µg/mL Gentamycin and 10% Fetal Bovine Serum (FBS) (Pan Biotech, cat. P30-3302).
CAL-27 cells were cultured in DMEM low glucose (Fisher Scientific, cat. 11966-025) supplemented with 10% FBS (Hyclone, cat. SH30109.03) and 1% Penicillin-Streptomycin.
For experiments in channels, cells were detached using Trypsin-0.25% EDTA (Fisher Scientific, cat 11570626), centrifuged and re-suspended in L-15 (Leibovitz Medium, Fisher Scientific, cat. 11540556) with the relevant concentrations: for the open-channels configuration, 100µL of a solution of 100,000 cells/mL is deposited on the PDMS motifs; for the closed-channels, a solution of 30,000,000 cells/mL is prepared and 20µL is injected in the chip reservoirs.

Gradient formation

A serum gradient is produced with two 1mL syringes prepared with L-15 0.1% BCS and 10% BCS with TRITC-dextran (20 kDa, Sigma, cat. 73766). The two solutions are flowed at 10µL/h in the chip inlets. After few minutes, a stable gradient is generated and can be visualized and quantified through imaging of the fluorescent dextran.

Immuno-fixation in micro-channels

Oxygen plasma treatment is done on the coverslip only, so that the chip can be reversibly bound to glass. Cells are seeded in the chambers as detailed above and are incubated during 12 hours at 37°C. The PDMS chamber is then immersed into a 3% PFA solution for 1h at room temperature. The chip is carefully removed with a scalpel and a tweezer. Cells are then permeabilized with Triton 0.5X for 3min and cells are incubated with staining agents. Phalloidin-AlexaFluor488 1:200 (Invitrogen, cat. A12379) was used to see actin and DAPI 1:1000 (Sigma, cat. 32670) to visualize nuclei.

Time-lapse Microscopy

For migration assays testing the bias and the computation of < p >, a standard phase-contrast microscope was used with a low magnification objective (10x, N.A. 0.4) with a time interval of 10min.
Nuclei deformation was imaged with an epifluorescence lamp (FluoArc Hg Lamp) coupled with a UV fluorescence filter. To do so, cells were incubated with DAPI (4µg/mL).
Actin bundles during migration were acquired with lifeAct-mcherry with the same epifluorescence microscope with a 40x oil objective (N.A. 1.3)
We visualized and tracked focal contacts with a stable NIH3T3 cell line expressing VASP-GFP.
Finally, keratin network in Cal-27 cells was acquired with a Nikon Spinning-Disk confocal microscope with a 60x oil objective (N.A. 1.4) using the Perfect Focus System.
Cell trajectories are generated by tracking the nucleus across time.

Cancer cell lines for keratin: K14 and fusion variant V7

Gene segments of wild type keratin 14 (K14) and the fusion variant 7 (V7) were amplified from oral cancer cells by nested PCR and sub-cloned into pAcGFP1-N1 vector (Clontech Lab., Madison. WI) as previously described (13). Because the main backbone of V7 is K14, K14 was utilized as the wild type control. The vector constructs were further transfected into CAL27 oral squamous cell line, purchased from the Bioresource Collection and Research Center (BCRC, Taiwan). The positive clones were enriched by cell sorting and maintained by G418 selection (700 μg/ml). The stable clones were generated by limiting dilution.

Polarity measurement

As a read-out for anisotropy in keratin (Fig. 2e-f), we defined the norm of the polarity $\vec{p}$ as following:

$$\left| \vec{p} \right|= \int_{0}^{S_{a}} I_{a}- \int_{0}^{S_{b}} I_{b}$$

where $I$ is the fluorescence intensity and $S$ the surface. The cell is divided in two regions $a$ and $b$ by its centre of mass.
